# FETCH enables fluorescent labeling of membrane proteins *in vivo* with spatiotemporal control in *Drosophila*

**DOI:** 10.1101/2025.01.31.635819

**Authors:** Kevin D. Rostam, Nicholas C. Morano, Kaushiki P. Menon, Davys H. Lopez, Lawrence Shapiro, Kai Zinn, Siqian Feng, Richard S. Mann

## Abstract

Fluorescent labeling approaches are crucial for elucidating protein function and dynamics. While enhancer trapping in *Drosophila* has been useful for the characterization of gene transcription, protein-specific visualization *in vivo* has been more elusive. To overcome these limitations, we developed Fluorescent Endogenous Tagging with a Covalent Hook (FETCH) to label cell surface proteins (CSPs) *in vivo* through a stable covalent bond mediated by the DogTag-DogCatcher peptide partner system^1^. FETCH leverages a spontaneous covalent isopeptide bond that forms between the 23-amino acid DogTag and the 15-kDa DogCatcher. Unlike most tags that work best at protein termini, DogTag is optimized for function in protein loops, expanding the range of sites that can be targeted in proteins. In FETCH, DogTag is introduced into extracellular loops of CSPs through genome engineering, enabling covalent bond formation with a genetically encoded DogCatcher-GFP fusion protein that can be secreted from a variety of cell types. We describe a flow cytometry-based platform for the identification of efficient DogTag insertion sites *in vitro* and demonstrate the ability to visualize both tagged DIP-α and Dpr10 *in vivo*, two immunoglobulin superfamily proteins that facilitate neuronal target recognition at *Drosophila* neuromuscular junctions and brain synapses. The versatility of FETCH enables fluorescent labeling with precise temporal and spatial control *in vivo*, enabling applications previously unfeasible.

## INTRODUCTION

The interaction network of DIPs (Dpr Interacting Proteins) and Dprs (Defective in Proboscis Extension Response^2^) was first characterized by an *in vitro* extracellular domain interaction assay for *Drosophila* cell surface proteins (CSPs)^3^. Further work has demonstrated that DIP-Dpr interactions facilitate neuronal target recognition and subsequent synaptogenesis in many tissues at all developmental stages^4–14^. The well-characterized interaction between DIP-α and Dpr10 is required for the stabilization of the neuromuscular junction (NMJ) at multiple developmental stages. The canonical view is that neuronal DIP-α interacts with Dpr10 molecules localized to the muscle cell membrane. In the larval neuromuscular system, genetic knockout of *DIP*-α or *dpr10* results in failed muscle innervation and mistargeting events by specific motor neuron axons^15–17^. Similarly, both *DIP-α* and *dpr10* are required for a subset of motor neurons to recognize and innervate their stereotyped muscles in the adult leg, despite their initial navigation to correct targets^18–20^.

While many proteins are amenable to direct fusions with fluorescent proteins, certain protein families such as the DIPs and Dprs are challenging to tag. The DIPs and Dprs consist of three and two immunoglobulin-like (Ig) domains, respectively, at their N-termini. The binding interface for DIP-Dpr interactions occurs between the most membrane-distal, N-terminal Ig1 domains of both proteins^4,21^. At their C-termini, both sets of proteins are tethered to the cell membrane via a glycosylphosphatidylinositol (GPI) anchor^22^. During translation and after entry into the endoplasmic reticulum, a C-terminal fragment of GPI-anchored proteins is cleaved at residues comprising the so-called ω-site, after which the mature protein is covalently bound to a GPI moiety and trafficked to the cell membrane^23^. These structural constraints make the termini of DIPs and Dprs resistant to modification, impeding the ability of standard labeling techniques that typically depend on tagging protein termini, such as affinity tags and fluorescent proteins, to visualize these molecules.

Due to these limitations, we developed a method based on DogTag-DogCatcher technology that we term “Fluorescent Endogenous Tagging with a Covalent Hook” (FETCH). This technology takes advantage of the loop-friendly DogTag-DogCatcher system, which enables spontaneous covalent bond formation between proteins^1^. The 23-amino acid DogTag peptide rapidly and spontaneously forms a covalent isopeptide bond with a specific binding partner, a 15 kDa protein denoted as DogCatcher^1^. The orthologous SpyTag-SpyCatcher pair, designed for ligation at protein termini, has previously been applied *in vivo* to tag transcription factor termini for chromatin immunoprecipitation (ChIP)^24^. However, the β strand conformation of SpyTag does not work well when inserted into protein loops. In contrast, DogTag has a β hairpin structure that is more compatible with insertion into protein loops, making it an optimal candidate for introduction into CSPs to mediate the covalent fusion with other proteins^1^.

We identify optimal DogTag insertion sites for DIP-α, Dpr10, and, for comparison, the transmembrane mouse CD8 (mCD8) protein, by using an easily scalable flow cytometry-based platform. CSPs with DogTag insertions are expressed on the surface of mammalian cells grown in culture, where ligation efficiency with DogCatcher can be quantitatively evaluated. We perform coupling reactions using recombinant expression of the most efficient DogTag variants of each CSP to biochemically confirm covalent bond formation. To study the localization of these proteins *in vivo*, we generated lines with optimized DogTag insertions and used DogCatcher-GFP to fluorescently label DIP-α, Dpr10, and mCD8 at *Drosophila* neuromuscular junctions. Finally, we describe a set of novel reagents that provide a secreted source of DogCatcher-GFP from a variety of cell types. The ability to rapidly identify optimal DogTag insertion sites in conjunction with the ease of genetic implementation positions FETCH as a versatile tool for protein-specific tagging and functional studies.

## RESULTS

### Overview of FETCH

In FETCH, a DogTag is introduced into the extracellular domains of CSPs of interest (Fig. 1A). DogCatcher, a 104 amino acid protein, is genetically encoded with a C-terminal fusion to a fluorescent protein and an N-terminal signal sequence to facilitate secretion *in vivo.* Our experiments *in vivo* use a heat shock-inducible promoter (*hsp70*) to allow temporal control of expression (Fig. 1B). In addition to DIP-α and Dpr10, we also introduce DogTag into mCD8, a mouse lymphocyte protein widely used as a membrane-localized marker of neuronal processes in *Drosophila*^25^. Vertebrate CD8 is also a member of the immunoglobulin superfamily (IgSF) class of cell surface proteins, and contains a single Ig-like domain in both CD8α and CD8β subunits^26^. Unlike DIPs and Dprs, CD8 is anchored to the membrane via a single pass transmembrane domain. To facilitate colocalization studies *in vivo*, we fused an mCherry fluorescent protein to mCD8α’s intracellular C-terminus (mCD8-mCherry). To identify suitable DogTag insertion sites in CSPs, we used a flow cytometry-based platform to quantify ligation efficiency between DogCatcher and a range of DogTag insertions into DIP-α, Dpr10, and mCD8-mCherry (Fig. 1C). This platform allows for the screening of large numbers of insertion sites in native membrane environments without necessitating recombinant protein production, making this approach readily scalable and adaptable to any CSP.

**Figure 1.**
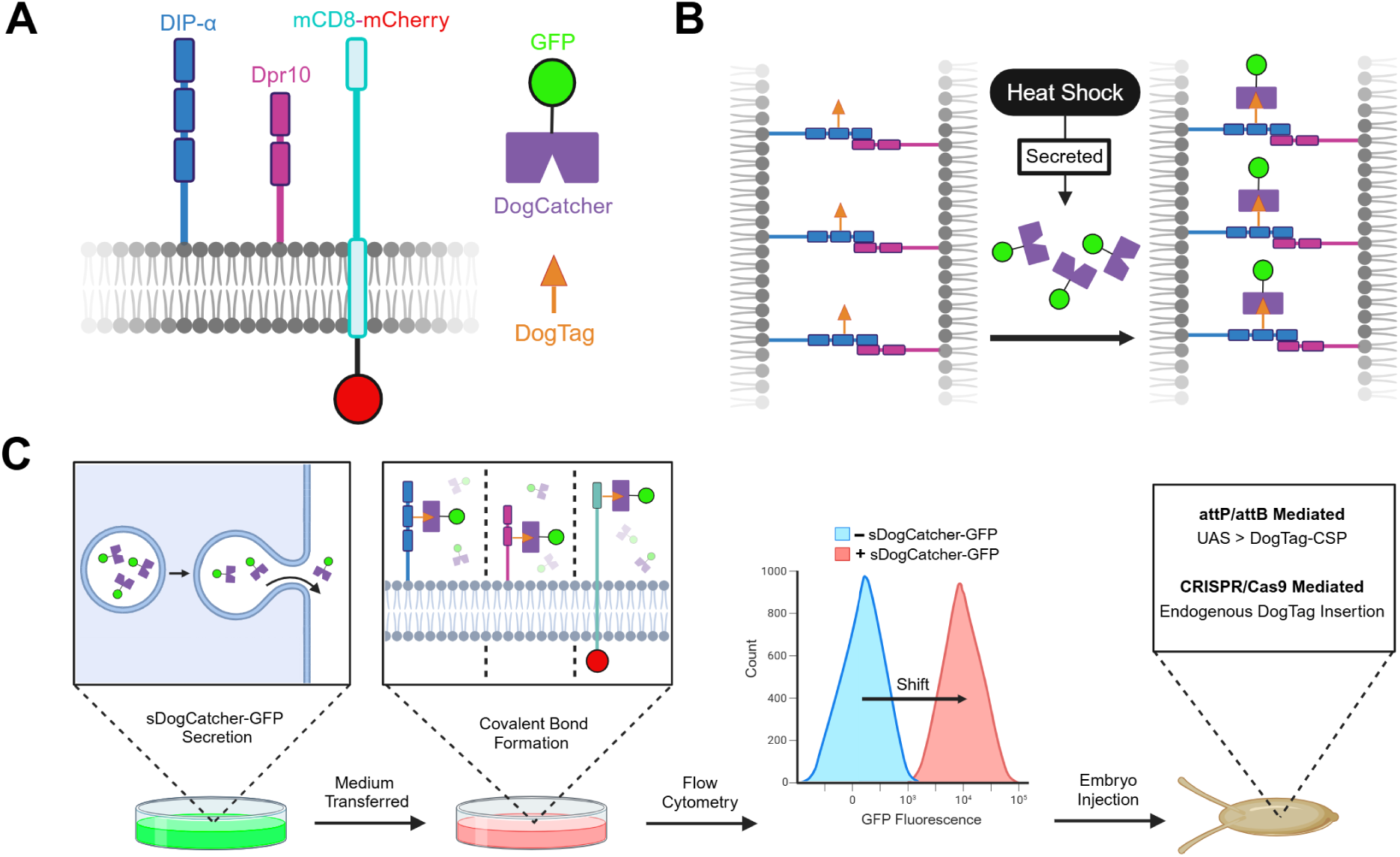
Overview of FETCH and flow cytometry platform for identifying successful DogTag insertion sites. **A.** Schematic of system components including the DogTag peptide, secreted DogCatcher fused to moxGFP (sDogCatcher-GFP), and IgSF cell surface proteins DIP-α, Dpr10, and mCD8. **B.** Cell surface proteins of interest are tagged with DogTag. An inducible heat shock promoter controls expression of secreted recombinant sDogCatcher-GFP protein *in vivo*. When in proximity to a DogTag, spontaneous covalent bond formation between DogTag and DogCatcher occurs, enabling protein-specific fluorescent tagging *in vivo*. **C.** Workflow for identifying optimal DogTag insertions for proteins of interest. Transfection of an sDogCatcher-GFP encoding plasmid into HEK293 cells results in fusion protein secretion into the supernatant. The medium is transferred to HEK293 cells expressing membrane-localized DogTag variants of DIP-α, Dpr10, or mCD8. Flow cytometry is used for quantitative comparison of DogTag candidates, which informs the choice of protein variants for further study and genome engineering.

### Flow cytometry quantifies the ligation efficiency of DogTag insertions into cell surface proteins

A variety of DogTag insertion sites were tested using our flow cytometry platform for all three proteins. For DIP-α and Dpr10, DogTag insertion into the N-terminal Ig1 domain of both proteins was avoided to prevent disrupting the binding interface required for their interaction (PDB: 6NRQ^4^) (Fig. 2E). Based on available crystal structures^4,26^, we targeted loops in the more membrane-proximal Ig domains of DIP-α (Ig2 and Ig3) and Dpr10 (Ig2), and in the sole Ig domain of the mCD8-mCherry fusion protein. We also inserted DogTags in the regions C-terminal to Ig3 for DIP-α or Ig2 for Dpr10, preceding the predicted sites of GPI linkage^20^. We refer to these regions as membrane linkers, which AlphaFold 3^27^ predicts to be unstructured (Fig. S1B and S1D). For mCD8-mCherry, the loops of its Ig domain, the unstructured membrane linker region, and a site immediately following its N-terminal signal peptide were targeted (Fig. S1F). In all cases, the 23 amino acid DogTag was flanked on both sides with a flexible five amino acid glycine-serine linker (see **Methods**).

**Figure 2.**
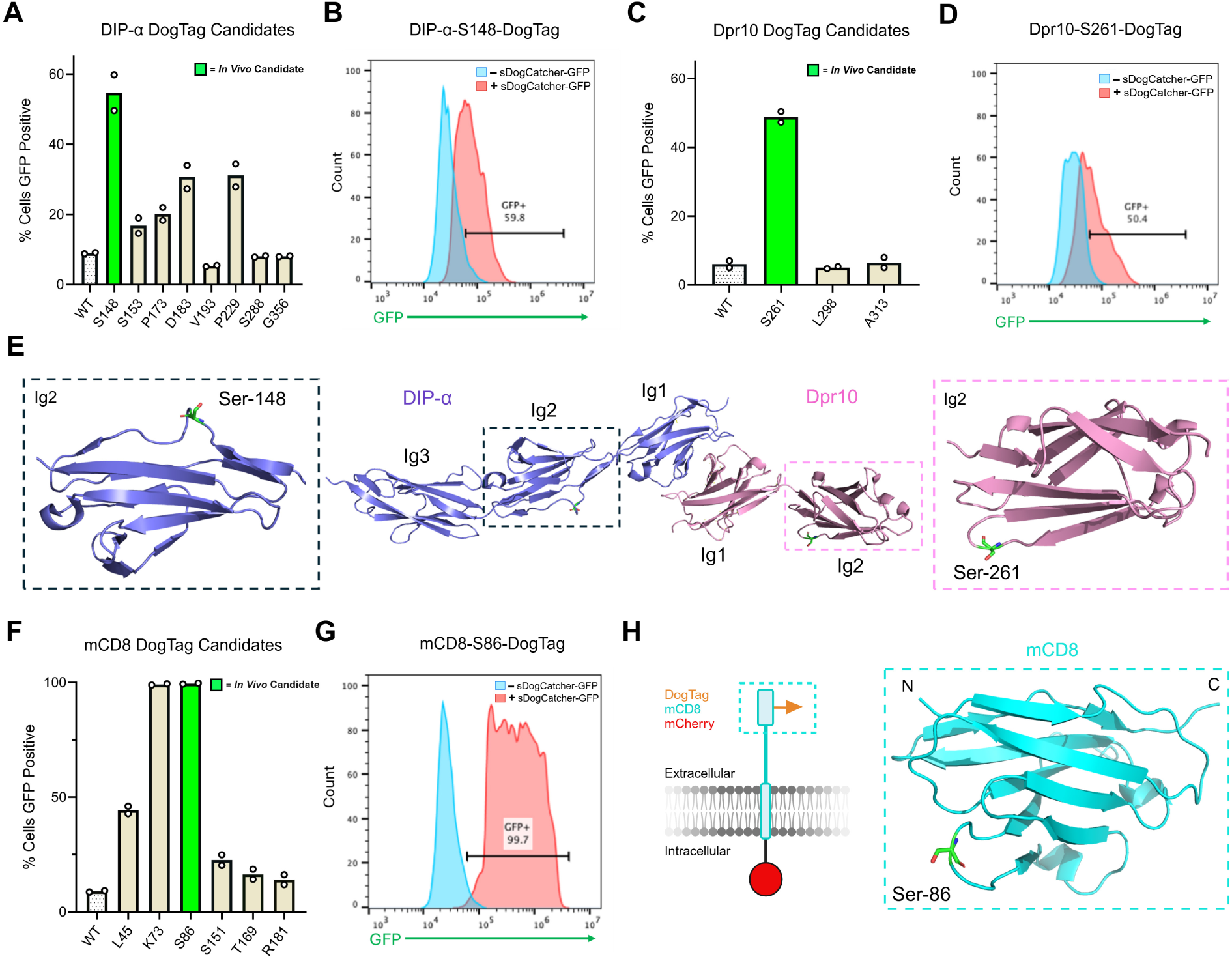
Flow cytometry quantifies the efficacy of covalent bond formation of DogTag insertion sites in DIP-α, Dpr10, and mCD8 *in vitro*. **A**, **C**, **F.** Bar graphs showing flow cytometry quantification of covalent bond formation between DogTag-CSPs and sDogCatcher-GFP. DogTag insertions were cloned into Ig2, Ig3, and the membrane linker region of DIP-α (**A**), Ig2 and the membrane linker region of Dpr10 (**C**), and the sole Ig domain and membrane linker region of mCD8 (**F**). DogTag insertions into loops after residues Ser148 for DIP-α (**A**), Ser261 for Dpr10 (**C**), and Ser86 for mCD8 (**F**) produced the largest fluorescent signal as detected by flow cytometry. DogTag was inserted immediately after the indicated residue numbers and flanked by flexible linkers in all constructs. Flow experiments were run in duplicate for all constructs. *WT* = Wildtype versions of respective cell surface proteins without DogTag. **B**, **D**, **G.** Histogram plots displaying change in fluorescent signal when supernatant containing sDogCatcher-GFP is incubated with HEK293 cells transfected with DIP-α-Ser148-DogTag (**B**), Dpr10-Ser261-DogTag (**D**), and mCD8-Ser86-DogTag (**G**) (red peaks), compared to cells incubated without sDogCatcher-GFP (blue peaks). An increase in fluorescence indicates covalent bond formation between the DogTag-CSP and sDogCatcher-GFP. FACS analysis of other DogTag-CSP variants and negative control WT proteins without DogTag are shown in Fig. S1. **E.** Crystal structure representation of the interaction between binding partners DIP-α and Dpr10 (PDBID: 6NRQ^4^). DogTag insertion sites at Ser148 and Ser261 for DIP-α and Dpr10, respectively, are indicated. **H.** Schematic of mCD8-DogTag-mCherry, where DogTag is inserted into the Ig domain and mCherry is fused to the truncated cytoplasmic domain. Crystal structure of mCD8’s Ig domain (PDBID: 2ATP^26^) with the DogTag insertion site at Ser86 is indicated.

A total of nine, three, and eight independent DogTag insertions were screened for DIP-α, Dpr10, and mCD8-mCherry, respectively. To assess covalent bond formation between DogTag-CSPs and DogCatcher, full length tagged CSP variants were transfected into HEK293 Freestyle cells. Expression levels were evaluated by monitoring cytoplasmic mCherry, which was expressed using the IRES system for DIP-α and Dpr10, and as a C-terminal fusion for mCD8-mCherry. We selected the cysteine-less moxGFP for fusion to DogCatcher, which is monomeric and engineered for use in the secretory pathway^28^. Recombinant DogCatcher-moxGFP, engineered with the secrecon signal peptide that drives robust secretion in mammalian cells^29^, was harvested from conditioned medium following transfection (see **Methods**). Secreted DogCatcher-moxGFP (here on referred to as sDogCatcher-GFP) was incubated with cells expressing DogTag-CSPs, which were then washed and then assessed for DogTag-DogCatcher ligation by flow cytometry with green fluorescence as the readout. For each CSP, multiple DogTag variants demonstrated coupling with sDogCatcher-GFP (Fig. S1A, S1C, and S1E). Among the DIP-α candidates, DogTag insertion into Ig2 after Ser148 achieved the largest fluorescent signal (Fig. 2A and 2B). Of the Dpr10 candidates, DogTag insertion into Ig2 after Ser261 produced the largest signal (Fig. 2C and 2D). For mCD8-mCherry, DogTag insertions into the Ig domain after Lys73 and Ser86 demonstrated the largest signals (Fig. 2F and 2G). Based on these results, DIP-α-S148-DogTag, Dpr10-S261-DogTag, and mCD8-S86-DogTag were identified as the most efficient variants and were selected for all subsequent biochemical and *in vivo* experiments (Fig. 2E and 2H). For simplicity, below we refer to these variants as DIP-α-DogTag, Dpr10-DogTag, and mCD8-DogTag-mCherry.

### Tagged loops achieve efficient protein-protein coupling *in vitro*

We compared the coupling efficiency of the loop-optimized DogTag-DogCatcher to SpyTag003-SpyCatcher003, a highly efficient orthologous system for ligation at protein termini^30^. For these experiments, the extracellular domains of DIP-α-DogTag (Ig1-Ig3), Dpr10-DogTag (Ig1-Ig2), and mCD8-DogTag-mCherry (sole Ig), were fused to a hemagglutinin (HA) signal peptide at their N-termini and SpyTag003 and 8xHis at their C-termini. Transfection of these expression constructs into Expi293 suspension cells resulted in protein secretion into medium, which was purified and concentrated for biochemistry. DogCatcher-GFP and SpyCatcher003 proteins were obtained by standard bacterial expression and purification. DIP-α-DogTag-SpyTag003-8xHis, Dpr10-DogTag-SpyTag003-8xHis, and mCD8-DogTag-SpyTag003-8xHis were independently incubated with either DogCatcher-GFP or SpyCatcher003 (Fig. 3A). Coupling reactions for all three CSP fragments resulted in prominent bands equivalent to the expected molecular weight of the ligation product for both DogTag-DogCatcher and SpyTag003-SpyCatcher003 coupling (Fig. 3B-D). Importantly, bands due to DogTag-mediated ligation in protein loops achieve comparable efficiency to SpyTag-mediated ligation at termini, revealing that DogTag-DogCatcher ligation is also highly efficient.

**Figure 3.**
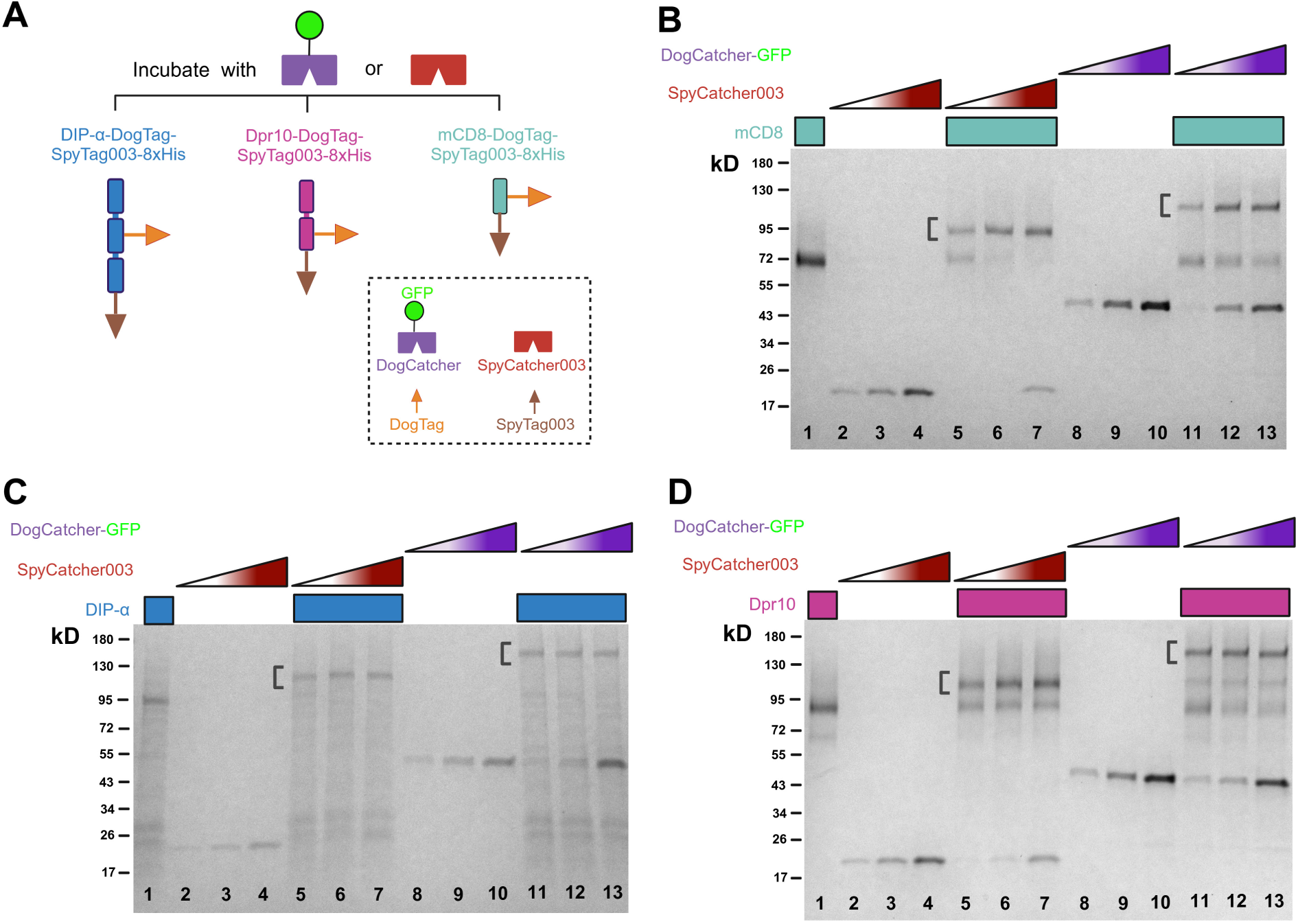
DogTag couples to DogCatcher with high efficiency in protein loops of DIP-α, Dpr10, and mCD8. **A.** Schematic showing soluble versions of DIP-α-DogTag, Dpr10-DogTag, and mCD8-DogTag that were generated by truncating their membrane linker regions, leaving Ig-like domains only. As a positive control for covalent bond formation, all three soluble protein variants were fused to a C-terminal SpyTag003, known to couple to SpyCatcher003 with high efficiency^30^. Soluble protein variants were incubated with increasing concentrations of DogCatcher-GFP or SpyCatcher003 in neutral buffer for 1 hour at room temperature. **B.** SDS-PAGE with approximately 5 pmols of mCD8-DogTag-SpyTag003-8xHis loaded in lanes 1, 5-7, and 11-13. DogCatcher-GFP or SpyCatcher003 amounts vary from half equimolar, equimolar, to twice equimolar relative to mCD8 in lanes 2-4, 5-7, 8-10, and 11-13, as indicated. Bands due to DogTag-DogCatcher coupling in lanes 11-13 have a similar efficiency to bands due to SpyTag003-SpyCatcher003 coupling in lanes 5-7. Molecular weight marker is shown on the left. **C.** SDS-PAGE with approximately 2.5 pmols of DIP-α-DogTag-SpyTag003-8xHis loaded in lanes 1, 5-7, and 11-13. DogCatcher-GFP or SpyCatcher003 amounts vary from half equimolar, equimolar, to twice equimolar relative to DIP-α in lanes 2-4, 5-7, 8-10, and 11-13, as indicated. Bands due to DogTag-DogCatcher coupling in lanes 11-13 have a similar efficiency to bands due to SpyTag003-SpyCatcher003 coupling in lanes 5-7. Molecular weight marker is shown on the left. **D.** SDS-PAGE with approximately 5 pmols of Dpr10-DogTag-SpyTag003-8xHis loaded in lanes 1, 5-7, and 11-13. DogCatcher-GFP or SpyCatcher003 amounts vary from half equimolar, equimolar, to twice equimolar relative to Dpr10 in lanes 2-4, 5-7, 8-10, and 11-13, as indicated. Bands due to DogTag-DogCatcher coupling in lanes 11-13 have a similar efficiency to bands due to SpyTag003-SpyCatcher003 coupling in lanes 5-7. Molecular weight marker is shown on the left. Brackets indicate protein bands resulting from DogTag-DogCatcher or SpyTag003-SpyCatcher003 ligation (**B, C, D**).

### FETCH enables labeling of CSPs at the larval neuromuscular junction *in vivo*

We generated a transgene in which expression of sDogCatcher-GFP is under the control of the heat shock-inducible *hsp70* promoter, allowing for temporally controlled expression in all cells (Fig. 4A). To boost the signal, we fused DogCatcher to 6 tandem copies of moxGFP (sDogCatcher-6xGFP), which resulted in a much stronger signal than a fusion with only 1 copy of moxGFP (Fig. S3). We also generated a transgene in which the *hsp70* promoter and sDogCatcher-6xGFP coding region were separated by a Flippase (Flp)-deletable FRT>STOP>FRT cassette, which we use below to restrict the secretion of sDogCatcher-6xGFP to specific cells. We verified secretion of DogCatcher *in vivo* by driving excision of the stop cassette in the *hsp70>STOP>sDogCatcher-6xGFP* transgene with Hedgehog-Gal4; UAS-Flp, allowing for heat shock-inducible expression only in the posterior compartment of the wing imaginal disc, and observed GFP fluorescence in the apical lumen of the anterior compartment (Fig. S2A).

**Figure 4.**
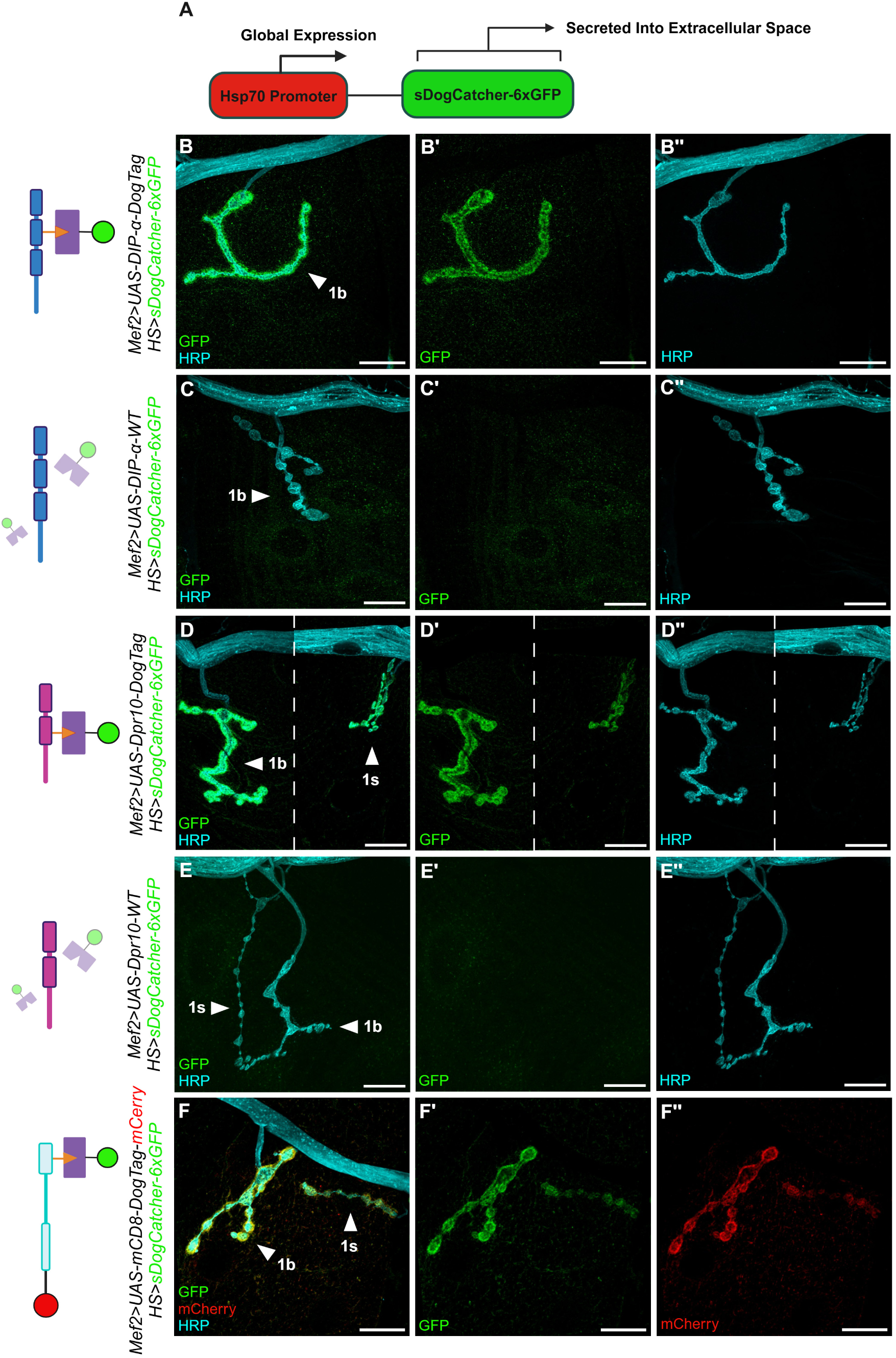
FETCH enables fluorescent labeling of DogTag-CSPs *in vivo*. **A.** Schematic demonstrating global expression of sDogCatcher-6xGFP, under the temporal control of the *hsp70* promoter. sDogCatcher-6xGFP secretion from all cells is achieved via standard heat shock. **B, B’, B”.** *Mef2-Gal4* drives expression of DIP-α-DogTag in third instar larval muscles, which localizes to the subsynaptic reticulum (SSR) of NMJs, as detected by ligation with global expression of sDogCatcher-6xGFP. A single confocal slice of Type 1b boutons at muscle 4 (m4) is shown as merged (**B**), GFP only (**B’**), and HRP only (**B”**) images. As observed previously^17^, the absence of the 1s bouton is a consequence of *cis* inhibition, when DIP-α is overexpressed in muscles. **C, C’, C”.** Mef2-Gal4 drives expression of DIP-α-WT in third instar larval muscles. Heat shock-induced sDogCatcher-6xGFP secretion occurs in the absence of a DogTag binding partner, with no detectable m4 SSR signal. A single confocal slice of Type 1b boutons at m4 is shown as merged (**C**), GFP only (**C’**), and HRP only (**C’’**) images. As observed previously^17^, the absence of the 1s bouton is a consequence of *cis* inhibition, when DIP-α is overexpressed in muscles. **D, D’, D”.** *Mef2-Gal4* drives expression of Dpr10-DogTag in third instar larval muscles, which localizes to the SSR of NMJs, as detected by ligation with global expression of secreted sDogCatcher-6xGFP. A single confocal slice of Type 1b and Type 1s boutons at m4 is shown as merged (**D**), GFP only (**D’**), and HRP only (**D”**) images. **E, E’, E”.** Mef2-Gal4 drives expression of Dpr10-WT in third instar larval muscles. Heat shock-induced sDogCatcher-6xGFP secretion occurs in the absence of a DogTag binding partner, with no detectable m4 SSR signal. A single confocal slice of Type 1b and Type 1s boutons at m4 is shown as merged (**E**), GFP only (**E’**), and HRP only (**E”**) images. **F, F’, F”.** Mef2-Gal4 drives expression of mCD8-DogTag-mCherry in third instar larval muscles, which localizes to the SSR of NMJs, as detected by ligation with global expression of sDogCatcher-6xGFP. A single confocal slice of Type 1b and Type 1s boutons is shown as merged (**F**), GFP only (**F’**), and mCherry only (**F”)** images. Arrowheads call out synaptic bouton types (**B-F**). Dashed lines denote separate focal planes within the same lateral plane (**D**). All scalebars = 20 µm.

Identical heat shock conditions in the absence of Flp expression produced no signal, suggesting that there is no transcriptional readthrough of the stop cassette (Fig. S2A). When mCD8-DogTag-mCherry is expressed in a subset of neurons in the larval ventral nerve cord, heat shock-induced global expression of sDogCatcher-6xGFP results in robust labeling only in the cells that express the tagged protein (Fig. S3A-D). Thus, these experiments confirm that DogTagged proteins can be recognized and bound *in vivo* by secreted sDogCatcher-6xGFP.

As a second test of FETCH, we used global heat shock-induced expression of sDogCatcher-6xGFP to label DIP-α-DogTag and Dpr10-DogTag when overexpressed at the larval neuromuscular junction (NMJ) using the muscle Gal4 driver, *Mef2-Gal4*. In both cases, we observe a robust GFP signal at the subsynaptic reticulum (SSR), invaginations of the muscle membrane that envelop 1b (big) and 1s (small) boutons, two types of glutamatergic motor neurons in the larval neuromuscular system^31^ (Fig. 4B and 4D). Notably, no signal was observed when untagged DIP-α-WT and Dpr10-WT were similarly overexpressed in the presence of global sDogCatcher-6xGFP expression (Fig. 4C and 4E). We also obtained robust labeling of mCD8-DogTag-mCherry in both 1b and 1s boutons when expressed with *Mef2-Gal4* (Fig. 4F). For these and all subsequent experiments using heat shock-induced expression of sDogCatcher-6xGFP, we found that a 30 minute heat shock, followed by a ∼12 hour chase prior to dissection, achieved an optimal signal over background, and without any aggregation of GFP (see **Supplemental Methods**). Together, these experiments demonstrate that overexpressed DogTag-CSPs can be successfully coupled to DogCatcher at neuromuscular junctions *in vivo*.

### Cell type-specific enhancers provide spatial control over DogCatcher-GFP expression

The experiments described above all rely on the *hsp70* promoter to drive temporally controlled, but ubiquitous expression of sDogCatcher-6xGFP. To provide spatial as well as temporal control over sDogCatcher-6xGFP secretion, we generated multiple *enhancer-Flp* transgenes to drive Flippase expression and restrict secretion to specific cell types (Fig. 5A). Based on previously described *enhancer-Gal4* transgenes, we chose enhancers with activity specific to immature myocytes *(Him-Flp*), mature myoblasts^32^ *(Mhc-Flp*), fat body^33^ *(3.1Lsp2-Flp*), glia^34^ *(Repo-Flp*), and both hemocytes and fat body^35^ (*Cg-Flp*). We confirmed their activity and cell-type specificity by staining pupae, using an *actin>STOP>GFP* transgene as our readout (Fig. S2B). In the larval NMJ, a glial source of sDogCatcher-6xGFP (*Repo-Flp; hsp70>STOP>sDogCatcher-6xGFP*) successfully labelled mCD8-DogTag-mCherry when expressed in the MNISN-1s glutamatergic motor neuron with DIP-α-Gal4 (Fig. 5B). Notably, in this case, labeling is observed in both the 1s bouton and the MNISN-1s axon, suggesting that the extracellular DogTag in mCD8 can be ligated to glia-derived sDogCatcher-6xGFP during trafficking in axons (Fig. 5B’). Labeling is undetectable in the absence of a Flp source (Fig. 5C).

**Figure 5.**
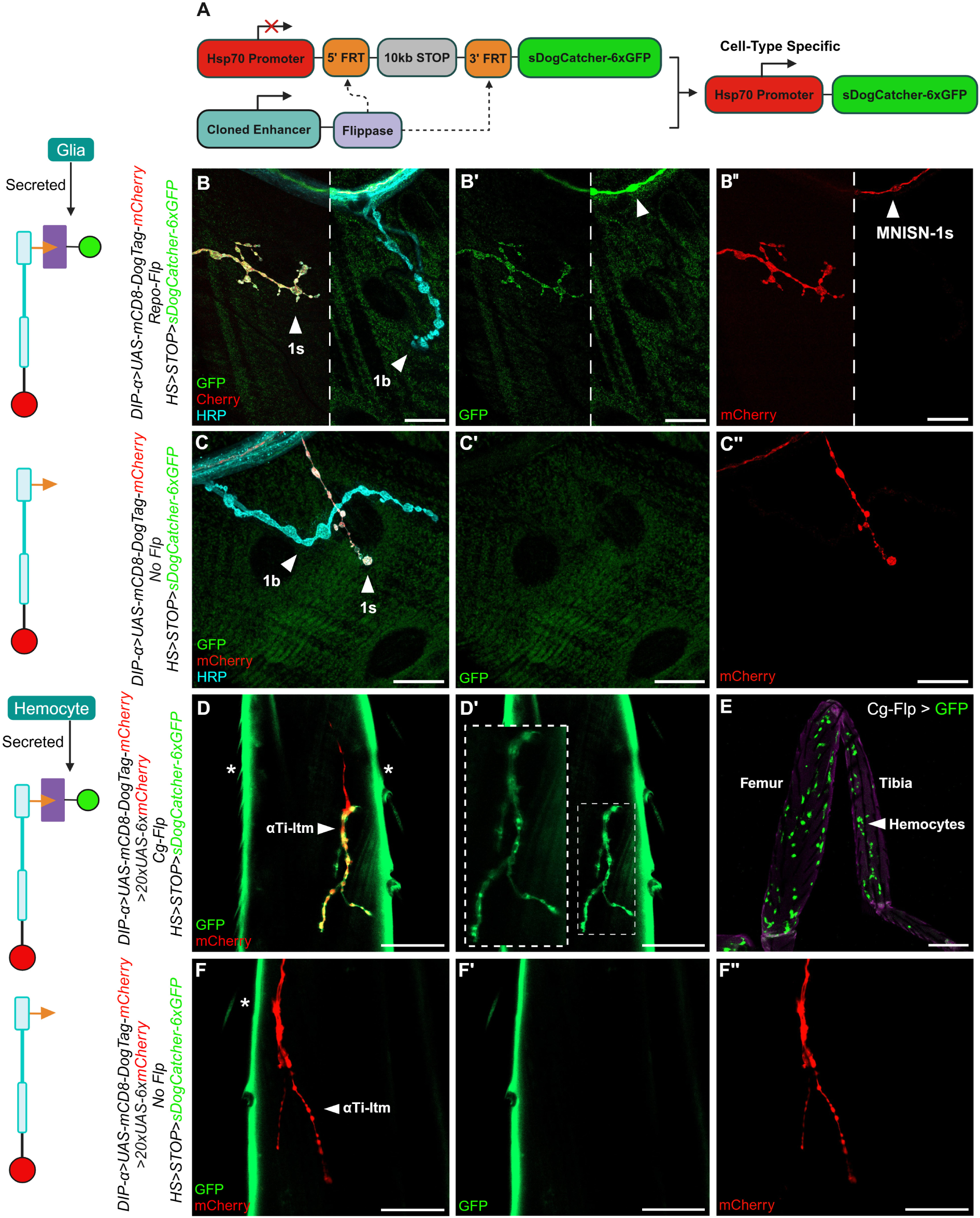
FETCH enables temporal and spatial control over fluorescent labeling in a cell type-specific manner. **A.** Schematic demonstrating a spatially restricted paradigm. Cell type-specific enhancers directly drive Flp expression, which excises an FRT-flanked stop cassette between *hsp70* and the sDogCatcher-6xGFP coding sequence, enabling temporal heat shock-inducible expression with cell type-specificity. **B, B’, B”.** *DIP-α-Gal4* drives neuronal expression of mCD8-DogTag-mCherry in the MNISN-1s neuron that innervates larval muscle 4 (m4). Note that GFP and mCherry signals are also observed in the nerve (arrowheads, **B’**-**B’’**). *Repo-Flp* deletes the FRT-flanked stop cassette to enable heat shock-induced sDogCatcher-6xGFP secretion from glial cells. A single confocal slice is shown as merged (**B**), GFP only (**B’**), and mCherry only (**B”**) images. **C, C’, C”.** *DIP-α-Gal4* drives neuronal expression of mCD8-DogTag-mCherry in the MNISN-1s neuron that innervates m4. Heat shock in the absence of a Flippase (Flp) source prevents stop cassette excision, transcriptionally terminating sDogCatcher-6xGFP expression. A single confocal slice is shown as merged (**C**), GFP only (**C’**), and mCherry only (**C”**). **D, D’.** *DIP-α-Gal4* drives neuronal expression of mCD8-DogTag-mCherry in the motor neuron targeting the long tendon muscle of the tibia in adult T1 legs (αTi-ltm). *DIP-α-Gal4* also drives cytoplasmic mCherry reporter expression. *Cg-Flp* deletes the FRT-flanked stop cassette to enable heat shock-induced sDogCatcher-6xGFP secretion from hemocytes. A single confocal slice is shown as merged (**D**) and GFP only (**D’**) images. Asterisks denote cuticle autofluorescence (**D**). Contrast of sDogCatcher-6xGFP localization is enhanced in inset (**D’**). **E.** Constitutive *Cg-Flp* expression deletes an FRT-flanked stop cassette between a constitutive actin promoter and Gal4 coding sequence, driving cytoplasmic GFP reporter expression in hemocytes in an adult T1 leg. Arrowhead calls out GFP-positive hemocytes. **F, F’, F”.** *DIP-α-Gal4* drives neuronal expression of mCD8-DogTag-mCherry in the αTi-ltm motor neuron in the adult T1 leg. *DIP-α-Gal4* also drives cytoplasmic mCherry reporter expression. Heat shock in the absence of a Flp source prevents stop cassette excision, transcriptionally terminating sDogCatcher-6xGFP expression. A single confocal slice is shown as merged (**F**), GFP only (**F’**), and mCherry only (**F”**). Asterisk denotes cuticle autofluorescence (**F**). Arrowheads call out synaptic bouton types (**B-D, F**). Dashed lines denote separate focal planes within the same lateral plane (**B**). Scalebars = 20 µm (**B**-**C)**, 25 µm (**D**, **F**), and 100 µm (**E**).

Next, we screened the ability of these cell type-restricted sources of sDogCatcher-6xGFP (*enhancer-Flp; hsp70>STOP>sDogCatcher-6xGFP*) to label mCD8-DogTag-mCherry in the adult leg NMJ. mCD8-DogTag-mCherry was expressed by DIP-α-Gal4, which is active in a subset of leg motor neurons^18^. Cytoplasmic mCherry reporter expression was included to unambiguously visualize the DIP-α-Gal4-expressing motor neurons. We observed the strongest signal at the leg NMJ when sDogCatcher-6xGFP was secreted from muscles and hemocytes, compared to fat body and glia (Fig. 5D and S4). While we present labeling of the motor neuron targeting the long tendon muscle of the tibia in adult T1 legs (αTi-ltm), we also observed strong labeling of all *DIP-α* expressing motor neurons in the tibia and femur. For labeling adult leg NMJs, *Cg*-*Flp*, which is restricted to hemocytes in the adult leg, emerged as the best non-neuronal and non-muscle source of sDogCatcher-6xGFP (Fig. 5E). Labeling of the leg NMJ is undetectable in the absence of a Flp source (Fig. 5F). Taken together, these experiments demonstrate that non-neuronal and non-muscle sources of sDogCatcher-6xGFP can successfully label extracellular domains of CSPs at neuromuscular junctions.

### FETCH enables visualization of DIP-α and Dpr10 expressed at endogenous levels

All of the above experiments rely on Gal4-UAS**-**driven expression of DogTagged CSPs, which can result in non-physiological expression levels and aberrant localization of these proteins. To determine if FETCH can label CSPs expressed at endogenous levels, CRISPR/Cas9 was used to introduce the DogTags described above for both DIP-α and Dpr10 at their native loci (Fig. 6A). We tested the ability of these proteins to be labeled at the larval NMJ, where innervation of the MNISN-1s motor neuron onto muscle 4 (m4) requires both DIP-α and Dpr10, expressed in the 1s motor neuron and SSR of m4, respectively^17^. Following heat shock-induced expression of sDogCatcher-6xGFP, we observe labeling of DIP-α-DogTag at the boutons of 1s neurons, but not 1b neurons, which do not express DIP-α (Fig. 6B). GFP labeling of tagged DIP-α is also observed in the MNISN-1s axon, suggesting that FETCH can label CSPs as they are trafficking in axons (Fig. 6B). Importantly, identical heat shock conditions in the absence of a DogTag-CSP demonstrate no promiscuous tagging in synaptic boutons of m4 and the MNISN-1s axon (Fig. 6D). For endogenously expressed Dpr10-DogTag, heat shock-induced expression of sDogCatcher-6xGFP labeled the SSRs enveloping both 1s and 1b boutons of m4 (Fig. 6C). Interestingly, the signal was stronger in the 1s boutons that have a less extensive SSR compared to 1b boutons, consistent with the idea that Dpr10 is stabilized by the presence of DIP-α, which is expressed in 1s, but not 1b motor neurons^21,36^. Notably, we also observed labeling of DIP-α-DogTag and Dpr10-DogTag at muscle 3 and muscle 2, which the MNISN-1s neuron also innervates.

**Figure 6.**
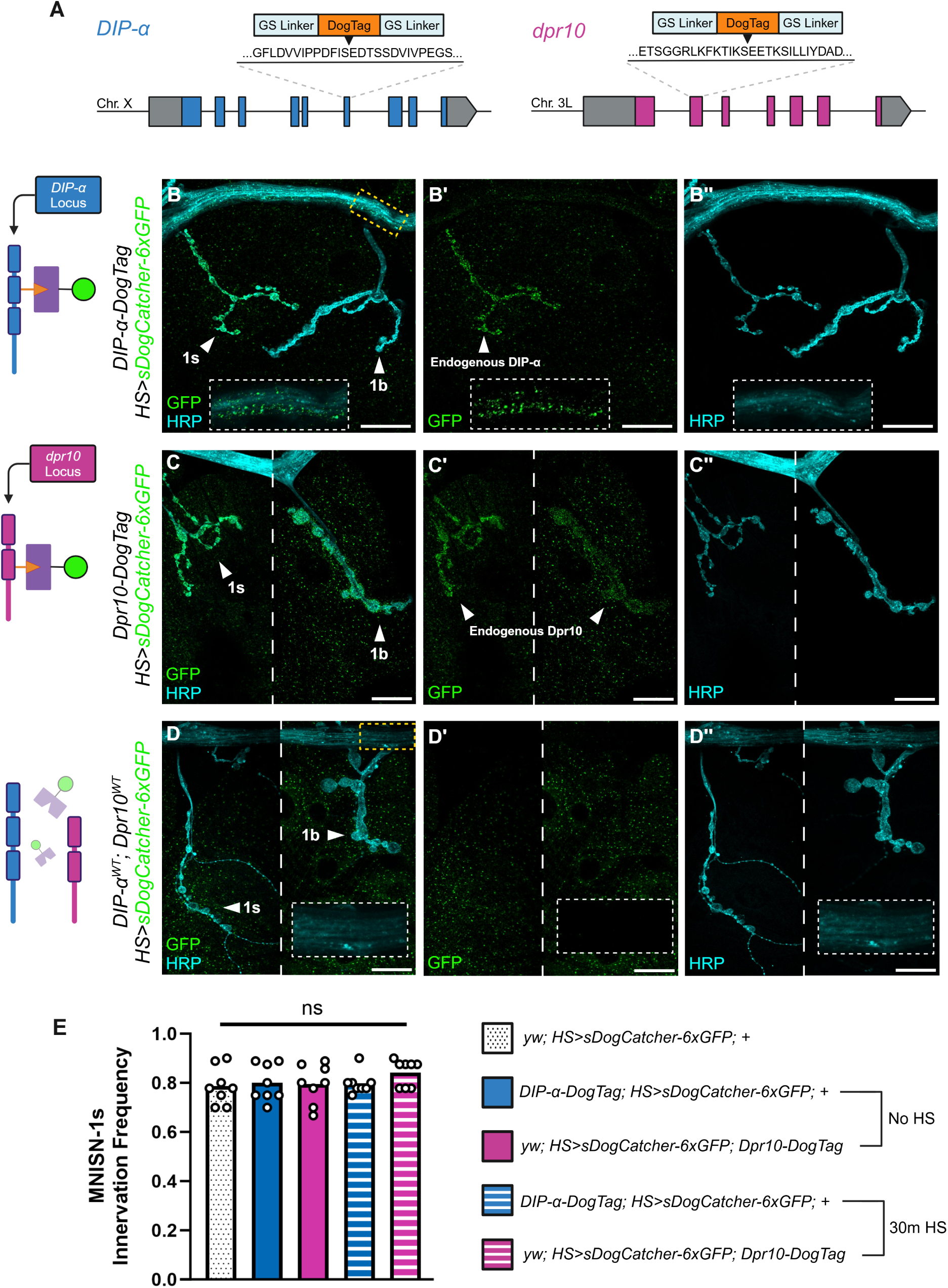
FETCH achieves fluorescent labeling of DIP-α and Dpr10 at endogenous expression levels. **A.** Schematic denoting locations of CRISPR-Cas9 mediated insertion of DogTag at the endogenous loci of *DIP-α* and *Dpr10*. Grey boxes denote 5’ and 3’ UTRs. Colored boxes denote exons. Black lines denote introns. **B, B’, B”.** Homozygous DIP-α-DogTag expressed at endogenous levels localizes to 1s boutons innervating third instar larval muscle 4 (m4), as detected by ligation to global expression of sDogCatcher-6xGFP. A single confocal slice is shown as merged (**B**), GFP only (**B’**), and HRP only (**B”**) images. White dashed insets correspond to yellow dashed box (**B**) and show active trafficking of DIP-α-DogTag in the MNISN-1s nerve. GFP and HRP signals in magnified insets are contrast enhanced. **C, C’, C”.** Homozygous Dpr10-DogTag expressed at endogenous levels localizes to the subsynaptic reticulum (SSR) surrounding 1s and 1b boutons innervating m4, as detected by ligation to heat shock-induced expression of sDogCatcher-6xGFP. A single confocal slice is shown as merged (**C**), GFP only (**C’**), and HRP only (**C”**) images. **D, D’, D”.** Heat shock-induced sDogCatcher-6xGFP expression with wildtype DIP-α and Dpr10 alleles does not result in promiscuous labeling in the absence of DogTag. White dashed insets correspond to yellow dashed box (**D**) and demonstrates no signal in the nerve. **E.** Quantitative comparison reveals no significant difference in MNISN-1s innervation frequency of third instar larval m4 in the presence of DIP-α-DogTag and Dpr10-DogTag alleles, both with and without ligation to heat shock-induced expression of sDogCatcher-6xGFP. Each datapoint represents the averaged % innervation frequency of multiple hemisegments in an individual animal. (Animals/Hemisegments) of genotypes in descending order of legend = (8/75), (8/67), (8/72), (8/75), (8/69). Abdominal hemisegments in A2-A6 were quantified for all animals. Genetic knockout of both DIP-α and Dpr10 results in total loss of MNISN-1s innervation of m4.^17^ *WT* = Wildtype proteins without DogTag. Arrowheads call out synaptic bouton types (**B-D**). Contrast enhancement of GFP and HRP signals in magnified insets (**B, D**) was performed consistently across samples and with the same confocal slices of respective figures. Scalebars = 20 µm (**B**-**D)**. Imaging and innervation frequency was determined in animals homozygous for all endogenous alleles and transgenes.

### DogTag-DogCatcher ligation does not disrupt the function of DIP-α and Dpr10

For FETCH to be a valuable tool, the covalent bond formed between the DogTag-CSP and sDogCatcher-6xGFP should not interfere with the function of the tagged protein of interest. The innervation frequency of MNISN-1s onto m4 has been extensively characterized as being highly sensitive to changes in the expression levels of both DIP-α and Dpr10^17^. We find that *DIP-α-DogTag* and *Dpr10-DogTag* homozygous animals, both with and without the expression of sDogCatcher-6xGFP, have a 1s innervation frequency comparable to wildtype animals, without tagged DIP-α or Dpr10 (Fig. 6E). These results provide strong evidence that the function of these CSPs is not compromised when bound to sDogCatcher-6xGFP. Based on these findings, we posit that DogTag-DogCatcher ligation can be used to fluorescently visualize CSPs without compromising biophysical interactions and functions *in vivo*.

## DISCUSSION

Fluorescent labeling approaches are critical for revealing protein localization and dynamics *in vivo*. FETCH overcomes several major limitations of existing techniques. Our development of a streamlined flow cytometry-based platform enables rapid identification and quantitative evaluation of DogTag insertion sites into membrane-localized CSPs *in vitro*, greatly accelerating the rate at which tagged CSP libraries can be screened. Current *in vivo* labeling approaches in *Drosophila*, such as split-fluorescent protein tags that reconstitute fluorescence^37^ or the recognition of epitope tags by nanobodies^38^, restrict the library of taggable proteins to those that can tolerate epitopes at their termini. These techniques also rely on the insertion of tandem repeats of tags to enhance detectability, demanding larger structural modifications to proteins that necessitates the generation of multiple tagged variants for each CSP, hindering adoption *in vivo*. FETCH facilitates fluorescent tagging within protein loops through a single DogTag insertion, with DogCatcher variants expressing a number of GFP copies tunable to expression levels of the tagged protein of interest. Since protein loops linking secondary structures are abundant in extracellular domains, FETCH can be harnessed to label both GPI-linked as well as transmembrane proteins in the immunoglobulin superfamily (IgSF) and other structural families of CSPs.

Surprisingly, in flow cytometry experiments, we observed that all DogTag insertions into the membrane linker regions (predicted to be unstructured) of all three CSPs (DIP-α-G356-DogTag, Dpr10-L298-DogTag, Dpr10-A313-DogTag, mCD8-S151-DogTag, mCD8-T169-DogTag, and mCD8-R181-DogTag), failed to produce notable increases in fluorescent signal, suggesting that they were unable to be productively bound by DogCatcher (Fig. S1A-F). Instead, the best insertion sites were located in small loops within otherwise well-structured Ig-like domains. It is plausible that the structure of DogTag, which adopts a β hairpin conformation, may not be stable in unstructured protein environments, thus hindering DogTag-DogCatcher isopeptide bond formation^1^. Based on our results, we suggest avoiding future DogTag insertions into regions predicted to be unstructured.

We observed that overexpression of both DIP-α-DogTag and DIP-α-WT in larval muscles resulted in a complete loss of MNISN-1s innervation of m4, consistent with previous observations^17^ and with the idea that interactions between DIP-α and Dpr10 in *cis* (in muscles) reduce the amount of endogenous Dpr10 available to interact with DIP-α in *trans*. Notably, *cis* inhibition by DIP-α and Dpr10 has also been observed in the adult leg neuromuscular system^20^. The ability of DIP-α-DogTag to cause *cis* inhibition is one indication that the tag does not compromise its ability to interact with Dpr10. Further, animals homozygous for the *DIP-α-DogTag* or *Dpr10-DogTag* alleles had normal MNISN-1s innervation frequency, both with and without ligation to sDogCatcher-6xGFP, suggesting that DogTag-DogCatcher chemistry and the covalent linking of fluorescent proteins in Ig2 does not interfere with the biophysical interaction required for their *in vivo* functions (Fig. 6E). Notably, when we expressed sDogCatcher-6xGFP globally to label CSPs, we are unable to distinguish whether covalent bond formation occurred as both proteins are moving through the secretory pathway or, alternatively, once the CSP is at the cell surface. In contrast, when sDogCatcher-6xGFP was expressed non-neuronally, it is more likely that labeling occurred at the cell surface. Regardless, although global expression of sDogCatcher-6xGFP worked well for DIP-α-DogTag and Dpr10-DogTag, the ability to have both temporal and spatial control of sDogCatcher-6xGFP expression *in vivo* may be important for functional studies requiring extracellular-specific labeling and/or labeling in a temporally controlled manner.

At endogenous expression levels of Dpr10-DogTag, we observed protein labeling at the SSR enveloping both 1s and 1b boutons of m4, with stronger labeling at 1s boutons (Fig. 6C). Interestingly, the m4 1b motor neuron does not express any of the known Dpr10 binding partners (DIP-α, DIP-β, nor DIP-λ)^21,36^. We suggest that our ability to visualize Dpr10-DogTag at the 1b NMJ is a consequence of the more extensive SSR membrane enveloping the boutons of 1b compared to 1s, and that the higher signal at the 1s boutons is a consequence of stabilization by a transsynaptic interaction with DIP-α^31,39^. Consistent with this idea, as with endogenously expressed Dpr10-DogTag, a stronger GFP signal at the 1s SSR was also observed when Dpr10-DogTag was overexpressed in larval muscles (Fig. 4D). In contrast, when mCD8-DogTag-mCherry, a molecule without known binding partners in *Drosophila*, was overexpressed in larval muscles and labeled by sDogCatcher-6xGFP, both the mCherry and GFP signals were stronger at the SSR of 1b compared to 1s (Fig. 4F), consistent with previous observations^40^. These differences in signal intensity, in which mCD8-DogTag-mCherry localizes less strongly to the 1s SSR compared to Dpr10-DogTag, reinforces the idea that Dpr10-DogTag localization, at both overexpressed and endogenous levels, is stabilized by its ability to interact with DIP-α.

Finally, we suggest that DogTag-DogCatcher technology has the potential for many additional *in vivo* applications beyond fluorescent labeling, in both invertebrate and vertebrate experimental systems. We envision that covalent interaction between DogTag and DogCatcher variants could also be used to force protein interactions between transsynaptic pairs, induce protein-specific degradation or proximity labeling, and visualize protein dynamics in live animals.

## Materials and Methods

### Cloning of DogTag and DogCatcher constructs

Synthesized Isoform A exon variants of DIP-α (UniProt ID: Q9W4R3) and Dpr10 (UniProt ID: Q9VT83) were used as template in the generation of DogTag DIP-Dpr variants (GenScript). pUAST-mCD8-GFP (Addgene #17746), encoding mouse CD8 subunit alpha, served as template for the generation of DogTag-mCD8 variants. DogTag candidates for each CSP were then subcloned into a bicistronic mammalian expression vector containing an internal ribosome entry site (IRES) for flow cytometry. Transfection into HEK293 cells resulted in the transcription of a single mRNA achieving DogTag-CSP expression from translation of the first cistron whereas translation of the second cistron would result in cytoplasmic mCherry reporter expression, serving as both a marker of transfection efficiency and a correlate of DogTag-CSP expression levels. For biochemistry, extracellular domain fragments of CSPs with DogTag insertions were adapted with a hemagglutinin (HA) signal peptide and C-terminal SpyTag003 and 8xHis-Tag through cloning into a CMV promoter-bearing mammalian expression vector. DogTag-CSPs and recombinant DogCatcher-moxGFP variants were cloned into plasmids containing attB sites for integration into the *Drosophila* genome through site-specific integration. pJ404-DogCatcher-sfGFP (Addgene #171930) served as template for the amplification of the 104 amino acid DogCatcher protein, and moxGFP (Addgene #68070) served as template for the amplification of moxGFP. DogTag was introduced into protein loops through overlap extension PCR. See **Supplemental Methods** for detailed cloning procedures.

### Tissue Culture and Transient Transfection

HEK293 Freestyle suspension cells were cultured in HEK Freestyle Media (Invitrogen, 12338018) and grown at 37° C in a humidified shaking platform incubator with 10% CO_2_. For transfection, cells were pelleted at 500x*g* and resuspended in fresh media. For small-scale (1 mL cells at 1×10^6^/mL) transient transfections performed in 24-well non-treated tissue culture plates, 10 μl of 293Fectin (ThermoFisher, 12347019) was added to 330 μL of Opti-MEM (ThermoFisher, 31985062), and incubated for 5 min at room temperature. 10 μl of transfection mixture was then added to 1000 ng of DNA, and incubated at room temperature for 30 min, after which it was added to HEK293 Freestyle cells in 24 well plates.

### Flow Cytometry Analysis of DogTag/DogCatcher Covalent Bond Formation

Flow cytometry titration assays were performed with cells expressing wildtype and DogTag variants of DIP-α, Dpr10, mCD8-mCherry, and sDogCatcher-GFP, transfected as described above. One day post transfection, cells were counted and diluted to 1×10^6^ cells/mL in 1xPBS 0.2% BSA, pH 7.4. Cells transfected with sDogCatcher-GFP were spun down at 500g for 2 minutes and supernatant was collected. Supernatant was then spun down at 2000g for 5 minutes to remove debris, and supernatant was collected again. 50 μL of DIP-α, Dpr10, and DogTag variant expressing cells were then added to 96 well plates (ThermoFisher, 262162) and combined with 50 μL of DogCatcher-GFP supernatant. 96 well plates were then incubated on an Elexa E5 Platform Shaker (New Brunswick Scientific) shaking at 250 rpm for 60 minutes at room temperature. Cells were then washed twice by centrifugation and removal of supernatant, and resuspending in 1xPBS 0.2% BSA. Cells were then analyzed by flow cytometry on a Novocyte Quanteon (Agilent). Gated live cells were sub-gated for high mCherry expression, and then sub-gated for GFP intensity. Analysis of flow data was done in FlowJo (FlowJo LLC).

### Transfection and Purification of Soluble Proteins

Expi293 suspension cells (ThermoFisher, A41249) were cultured in Expi293 medium (ThermoFisher, A1435101) and grown at 37° C in a humidified shaking platform incubator at 130R PM with 8% CO2. For transfection, cells were introduced into prewarmed Expi293 medium at a concentration of 3×10^6^ cells/mL. In two separate tubes, 50 µL of Opti-MEM (ThermoFisher, 31985062) per 1 mL transfection were prepared. 1 µg of plasmid DNA per 1 mL transfection was added to the first tube. 2.7 µL ExpiFectamine 293 Reagent (ThermoFisher, A14525) per 1 mL transfection was added to the second tube and mixed before incubation at room temperature for 3 minutes. DNA and ExpiFectamine tube contents were mixed and incubated at room temperature for 5 minutes. The mixture was then added to Expi293 cells and shaken at aforementioned conditions. 16-20 hours later, ExpiFectamine 293 Transfection Enhancers 1 and 2 (ThermoFisher, A14525) were added to cells at 5 µL and 50 µL per 1 mL of transfection, respectively. 96 hours later, cells were pelleted at 4000 RPM for 15 minutes at 4° C. Supernatant was loaded through a 1mL HisTrap FF column (Cytiva Life Sciences, 17531901) at 0.5mL/min using a peristaltic pump. The column was washed with 10 mL of 20mM imidazole (ThermoFisher, A10221) in 1xPBS. Protein was eluted with 5 mL of 250mM imidazole in 1xPBS.

### Coupling Reactions

Purified proteins were kept at −80° C for long term storage. Small aliquots were kept at 4° C for immediate experimental use. For Dpr10-DogTag-SpyTag003-8xHis and mCD8-DogTag-SpyTag003-8xHis, protein amount was constant at 5 pmols across all ligation experiments. For both DogCatcher-GFP and SpyCatcher003, protein amounts of 2.5 pmols, 5 pmols, and 10 pmols were used in independent coupling reactions with 5 pmols of the aforementioned tagged extracellular domain fragments. For DIP-α-DogTag-SpyTag003-8xHis, protein amount was constant at 2.5 pmols across these ligation experiments. DogCatcher-GFP and SpyCatcher003 protein amounts of 1.25 pmols, 2.5 pmols, and 5 pmols were used in independent coupling reactions with 2.5 pmols of this tagged extracellular domain fragment. For all coupling reactions, proteins were incubated together in 8-strip PCR tubes at room temperature in 1x PBS (pH 7.4) for 1 hour in a total volume of 10 µL. For all experiments, proteins were also ran alone at aforementioned amounts and subjected to conditions identical to those for coupling. Afterwards, 3.33 µL 4xSDS-PAGE loading dye with 10% β-mercaptoethanol was added to samples and incubated for 10 minutes at 98° C prior to SDS-PAGE.

### SDS-PAGE Analysis

Purified protein was concentrated further through 10 kDa MWCO Ultra Centrifugal Filter (Millipore, UFC9010). 4xSDS-PAGE loading dye with 10% β-mercaptoethanol was added to samples before a 10 minute incubation at 98° C. Proteins were run on 12-well precast protein gels (Bio-Rad, 4561085) with BSA standards (ThermoFisher, 23208) to obtain protein concentrations. For all coupling reactions, proteins were incubated in a total volume of 10 µL. 3.33 µL 4xSDS-PAGE loading dye with 10% β-mercaptoethanol was added to samples and incubated for 10 minutes at 98° C. Proteins were then run on 15-well precast protein gels (Bio-Rad, 4651086). Gels were stained with Coomassie InstantBlue (Abcam, ab119211) and imaged with a ChemiDoc XRS+ molecular imager (Bio-Rad).

### Transgenesis

All plasmids for site-specific recombination and introduction into the *Drosophila* genome carried *attB* sites. *Enhancer-Flp* constructs were injected into *attP40w-* flies (Rainbow Transgenic Flies). UAS expression constructs of tagged and wildtype CSPs (DIP-α, Dpr10, and mCD8-mCherry) were injected into *attP2w-* flies (Rainbow Transgenic Flies). sDogCatcher-GFP constructs were injected into *attP1w-* flies (Rainbow Transgenic Flies). See **Supplemental Methods** for enhancer isolation and detailed transgene details.

### Generation of *DIP-α-DogTag* and *Dpr10-DogTag* CRISPR Alleles

We chose protospacer sequences in the exon regions of DIP-α and Dpr10 to insert DogTag after Ser148 and Ser261, respectively. DogTag insertions were flanked with GGGGS for DIP***-***α and GGGS for Dpr10. High score protospacer sequences were chosen on http://crispr.dfci.harvard.edu/SSC/. We cloned independent protospacers into pCFD3-dU6:3gRNA (Addgene #49410) and co-injected the plasmid and single-stranded repair template (Integrated DNA Technologies) into CAS0001 flies (Rainbow Transgenic Flies). Injected embryos were crossed to balancer lines, and F1 individuals were PCR screened for successful mutagenesis. Stable stocks were established from individuals demonstrating successful DogTag insertion and correct genomic repair. sgRNA and ssODN sequences are below. Detailed protocols available upon request.

DIP-α-DogTag sgRNA: TCCGCCGGACTTCATCAGCG

DIP-α-DogTag ssODN (5’ to 3’): CGCACGGAACTGCCCTCCGGCACAATCACATCGGATGAGGTGTCCTCGGAACCACCGCCGCCTTTCGGCGGTATCGGTTCATTGGTGATATAATGTTTACCATCGGTGAATTCGTATGTAGCCGGAATATCCGAGCCGCCTCCTCCGCTGATGAAGTCCGGCGGAATCACGACGTCCAGGAAGCCAATCTACAATTACC

Dpr10-DogTag sgRNA: ATTCAAGACCATCAAGTCGG

Dpr10-DogTag ssODN (5’ to 3’): CAGAGTGCAGGAGATCCGCATCGTAAATGAGCAAAATGGATTTGGTCTCCTCGGAACCGCCGCCTTTCGGCGGTATCGGTTCATTGGTGATATAATGTTTACCATCGGTGAATTCGTATGTAGCCGGAATATCCGAGCCTCCTCCCGACTTGATGGTCTTGAATTTCAGTCGGCCGCCCGAAGTTTCCTCGCTGAGCACC

### Dissections

Larval fillets were dissected in cold 1xPBS and fixed for 25 minutes with 4% PFA in PBS. Fillets were washed 4x for 10 minutes in 1xPBS, and then 3x for 10 minutes in cold PBST-BSA (0.05% Triton, 0.1% BSA) on a nutator at room temperature. PBST-BSA was replaced with 5% heat-inactivated normal goat serum (Gibco, 16210064) in PBST-BSA and blocked overnight on a nutator at 4° C. Alexa Fluor 647-conjugated goat anti-horseradish peroxidase (Jackson ImmunoResearch, 123605021; 1:100) was incubated with filets in 5% HI-NGS in PBST-BSA overnight on a nutator at 4° C. Fillets were then washed 4x for 10 minutes with PBST-BSA and 2x for 10 minutes with 1xPBS on nutator at room temperature. Samples were then placed ventral side downwards and mounted in Vectashield (Vector Laboratories, H100010). For adult leg dissections, flies were immersed in 70% ethanol for 30 seconds and rinsed 3x in PBST (0.3% Triton). Abdomen and heads were removed and legs still attached to thoracic segments were fixed with 4% PFA-PBST overnight on a nutator at 4° C. Then, legs and thorax were washed 5x in PBST and stored overnight in 80% mounting medium (Vector Laboratories, H100010) in 1xPBS overnight at 4° C. T1 legs were then removed from thoracic segments and mounted with dental wax on the corners of coverslips to accommodate leg thickness.

### Microscopy

A Zeiss Axio Imager 2 based LSM800 confocal laser scanning microscope was used for all microscopy experiments. A 63x/1.4 NA oil-immersion objective was used for imaging single slices for larval and adult neuromuscular junctions. A 25x/0.8 NA oil-immersion objective was used for imaging larval VNCs with a step size of 0.5 µm. A 10x/0.3 NA non-confocal objective was used for imaging wing imaginal discs and pupae with step sizes of 0.5 µm and 1.0 µm, respectively. Laser power and gain settings were identical across samples in each set of experiments. Post image processing was identical across samples in each set of experiments and performed with ImageJ software.

### Statistical Analysis

Analysis of innervation frequency was performed with GraphPad Prism 10.1.0 software (San Diego, CA, USA). MNISN-1s innervation of third instar larval muscle 4 was quantified as either present (1) or absent (0), with values across hemisegments averaged per animal. Statistical analysis for significance between conditions was evaluated with an unpaired, non-parametric Mann-Whitney U test.

## ACKNOWLEDGEMENTS

This work was supported by NIH grants R01 NS070644 and R35 GM118336 to R.S.M., and NSF grant IOS-2321481 to K.Z. and L.S. We thank William J. Glassford for input in designing the Dpr10-DogTag CRISPR allele. We thank Michael Anaya and Annie W. Lam for assistance with mammalian expression and purification of proteins for biochemistry experiments. We thank Frank Schnorrer for providing details for the *Him* enhancer, Gary Struhl for sharing *hh-Gal4* reagents and stop cassette DNA, Alina P. Sergeeva for discussions about DogTag insertions and Jordan Becker for assisting with flow cytometry transfections. We thank all members of the Mann lab for discussions. We acknowledge the support of Columbia University’s Summer Undergraduate Research Fellowship (SURF) program for supporting K.D.R. in the lab of R.S.M. in 2023 and the Amgen Foundation for supporting K.D.R. as an Amgen Scholar at the California Institute of Technology in the lab of K.Z. in 2024.

## AUTHOR CONTRIBUTIONS

K.D.R., S.F., and R.S.M. conceptualized the project. K.D.R. conducted flow cytometry experiments with N.C.M., who designed and developed the flow cytometry platform. K.D.R. conducted biochemistry experiments at the California Institute of Technology with input from K.Z. K.D.R. generated CRISPR/Cas9 alleles and transgenes. K.D.R. conducted larval and adult NMJ experiments with input from K.P.M. K.D.R. and D.H.L. performed pupal screening of *enhancer-Flp* constructs. R.S.M. supervised experiments and provided funding. K.D.R. wrote the manuscript. K.D.R., N.C.M., K.P.M., and R.S.M. edited and revised the manuscript. D.H.L., K.Z., and S.F. commented on the manuscript, and all authors approved the final draft.

---

**Figure S1.**
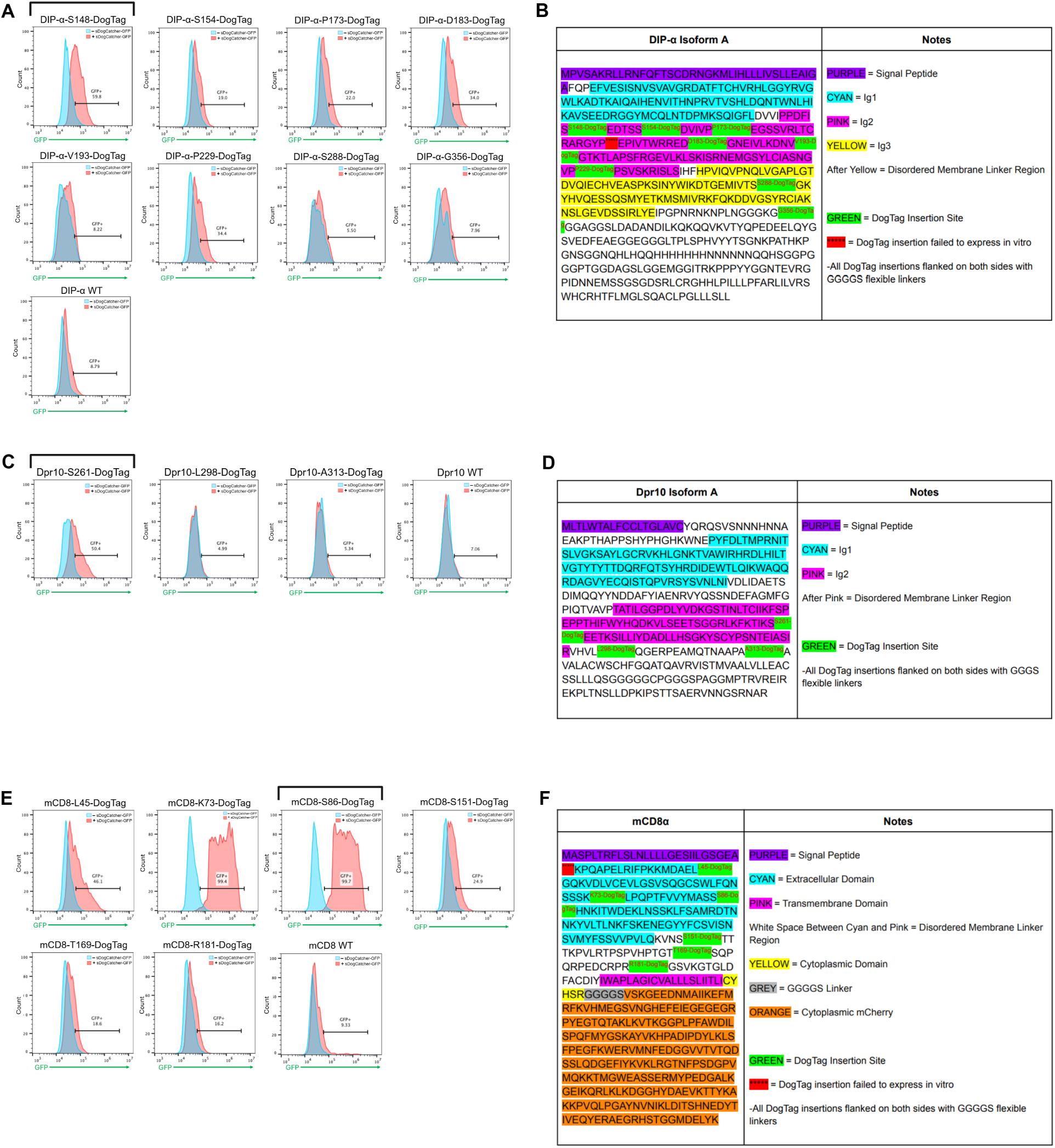
Full flow cytometry dataset and annotation of all DogTag insertions into extracellular domains of DIP-α, Dpr10, and mCD8 *in vitro*. **A.** Nine DogTag insertions were cloned into Ig2, Ig3, or the membrane linker region of DIP-α, one of which failed to express. As such, there are eight histogram plots for DogTag variants in addition to a wildtype control. Flow experiments were run in duplicates for all constructs. Black bracket denotes optimal variant used in subsequent experiments. *WT* = Wildtype DIP-α without DogTag. **B.** Primary structure annotation of DIP-α. Tested DogTag insertion sites and protein domains are indicated. **C.** Three DogTag insertions were cloned into Ig2 or the membrane linker region of Dpr10. There is an additional histogram plot for a wildtype control. Flow experiments were run in duplicates for all constructs. Black bracket denotes optimal variant used in subsequent experiments. *WT* = Wildtype Dpr10 without DogTag. **D.** Primary structure annotation of Dpr10. Tested DogTag insertion sites and protein domains are indicated. **E.** Six DogTag insertions were cloned into the sole immunoglobulin-like domain or membrane linker region of mCD8, with an seventh insertion after the native signal peptide failing to express. As such, there are six histogram plots for DogTag variants in addition to a wildtype control. Flow experiments were run in duplicates for all constructs. Black bracket denotes optimal variant used in subsequent experiments. *WT* = Wildtype mCD8-mCherry without DogTag. **F.** Primary structure annotation of mCD8-mCherry. Tested DogTag insertion sites and protein domains are indicated.

**Figure S2.**
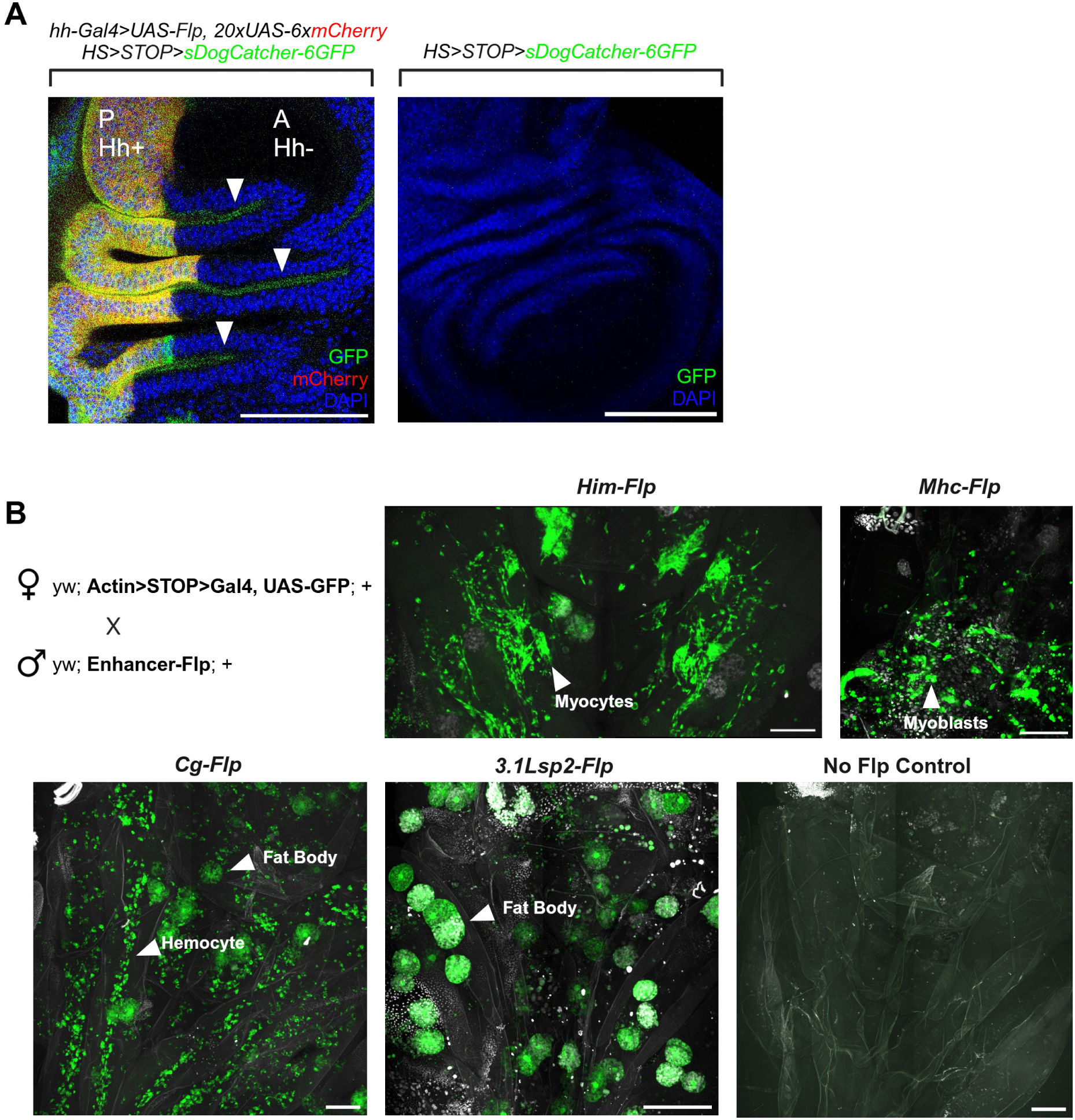
Functional validation of DogCatcher secretion, transcriptional termination by stop cassette, and enhancer-Flp constructs. **A.** Hedgehog-Gal4 expression, isolated to the posterior compartment of the wing disc, drives expression of cytoplasmic mCherry. Hedgehog-Gal4 also drives Flippase (Flp), flipping out an FRT-flanked stop cassette between *hsp70* and the coding region of sDogCatcher-6xGFP. Heat shock demonstrates DogCatcher secretion through the apical end of the wing disc into the lumen. Samples with identical heat shock conditions devoid of Flp expression demonstrates no DogCatcher transcription or secretion. Arrowheads indicate luminal sDogCatcher-6xGFP. *Hh* = hedgehog. *P* = wing disc posterior compartment. *A* = wing disc anterior compartment. **B.** A variety of enhancer fragments were isolated from the *Drosophila* genome and cloned to drive Flippase1 (Flp) expression. *Enhancer-Flp* flies were crossed with animals possessing an FRT-flanked stop cassette between a constitutive actin promoter and a Gal4 coding region. Gal4 expression then drives cytoplasmic GFP in a cell-type specific manner. *Mhc* = 2.4kb regulatory region of *myosin heavy chain*. *Him* = 4.3kb regulatory region of *holes in muscles*. *Cg* = 2.7kb regulatory region between *Col4a1* and *vkg*. *3.1Lsp2* = 3.1kb regulatory region of *larval serum protein 2*. Arrowheads call out cell-type specific Flp expression. Scalebars = 100 µm in **A** and **B**.

**Figure S3.**
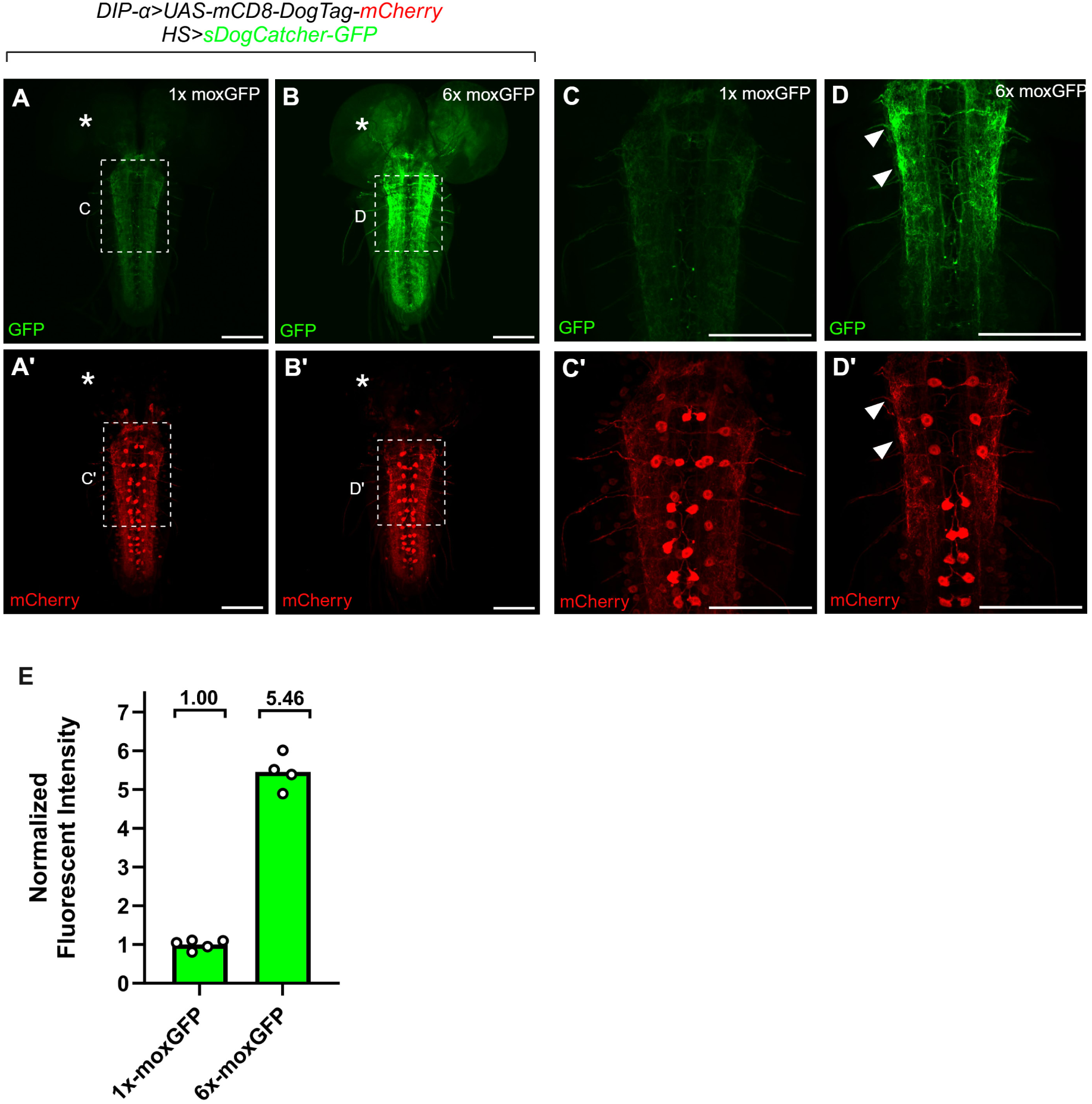
mCherry-moxGFP colocalization from ligation of mCD8-DogTag-mCherry to sDogCatcher with and without tandem repeats of moxGFP. **A-D.** DIP-α-Gal4 driving mCD8-DogTag-mCherry expression in larval ventral nerve chords (VNC) with sDogCatcher-GFP expression under temporal control of *hsp70*. One hour heat shock was followed by two hour chase with sDogCatcher fused to either one copy of moxGFP (**A, C**) or six tandem repeats of moxGFP (**B, D**). Arrowheads call out colocalization. Asterisks call out lack of mCD8-DogTag-mCherry expression and absence of GFP signal in the optic lobe. Scalebars = 100 µm. **E.** Quantitative comparison of fluorescent intensity between one and six copies of moxGFP fused to sDogCatcher after coupling to mCD8-DogTag-mCherry in the larval VNC. Region of interest (ROI) = VNC without optic lobes. Fluorescent intensity of maximum intensity projections were quantified with ImageJ and normalized to 1.00.

**Figure S4.**
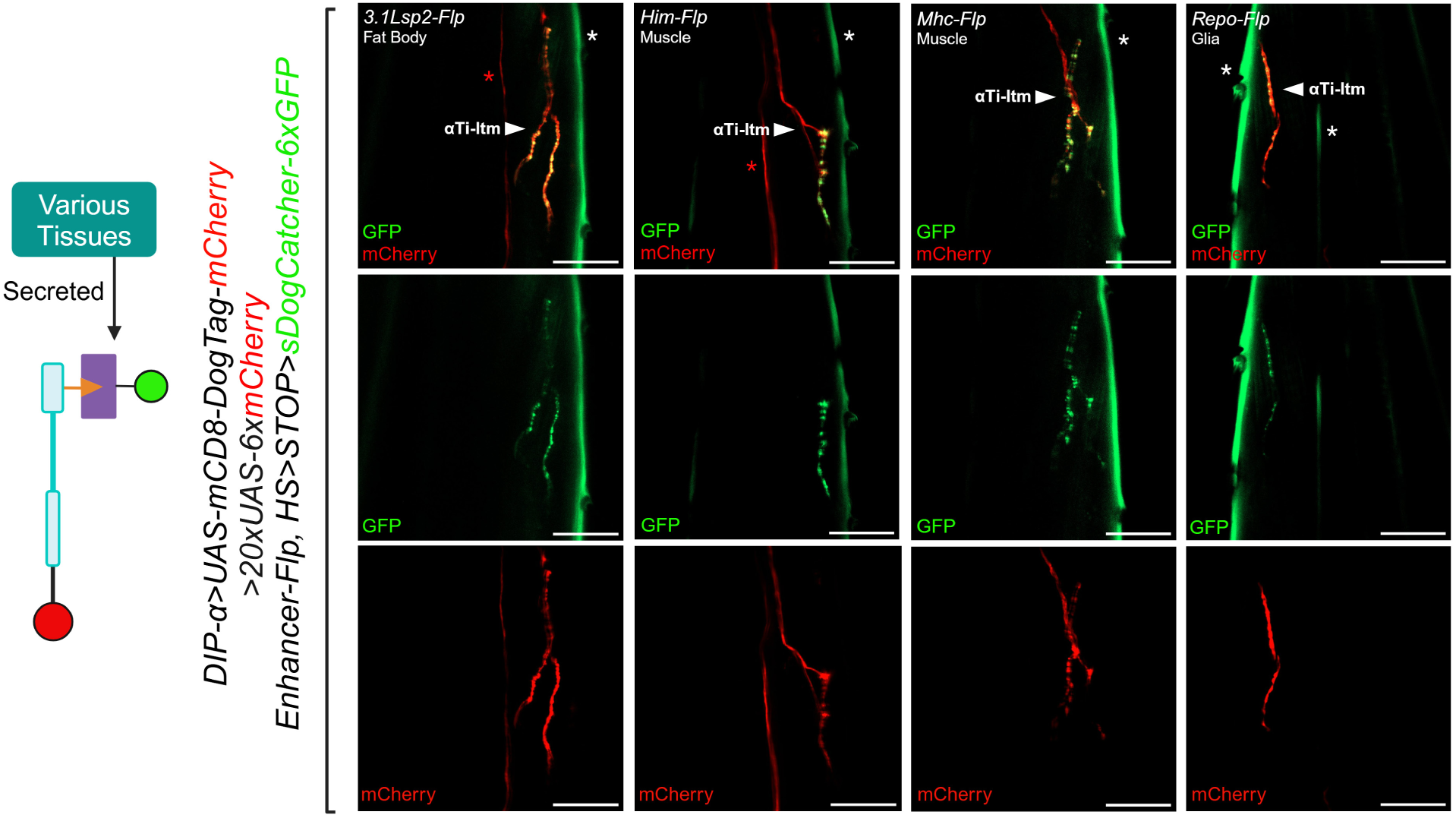
Screening of *enhancer-Flp* transgenes through spatially restricted labeling of adult leg neuromuscular junctions. DIP-α-Gal4 drives neuronal expression of mCD8-DogTag-mCherry in motor neuron targeting the long tendon muscle of the tibia in adult T1 legs (αTi-ltm). DIP-α-Gal4 also drives cytoplasmic mCherry reporter expression. Various Flp constructs excise the FRT-flanked stop cassette to enable heat shock-induced sDogCatcher-6xGFP expression from denoted tissues. Single slices presented as merged, GFP only, and mCherry only images. White asterisks denote cuticle autofluorescence. Red asterisks denote cytoplasmic mCherry reporter expression in motor neuron axons targeting the tarsal depressor muscle of the tibia in adult T1 legs. *WT* = Wildtype proteins without DogTag. Arrowheads call out synaptic bouton types. Scalebars = 25 µm.

